# A Comparative Evaluation of Molecular Connectivity and Covariance Approaches

**DOI:** 10.64898/2025.12.17.694867

**Authors:** Murray B Reed, Samantha Graf, Matej Murgaš, Benjamin Eggerstorfer, Christian Milz, Pia Falb, Elisa Briem, Alexandra Mayerweg, Gabriel Schlosser, Lukas Nics, Godber Mathis Godbersen, Sazan Rasul, Marcus Hacker, Andreas Hahn, Rupert Lanzenberger

## Abstract

Advances in high-temporal-resolution functional positron emission tomography (fPET) now enable the assessment of metabolic associations between brain regions, providing a molecular complement to functional connectivity derived from fMRI. However, the distinction between molecular connectivity (MC) and covariance (mCov) remains conceptually and methodologically inconsistent across studies. This work systematically compares major analytical approaches for MC and mCov to clarify their assumptions, dependencies, and interpretational boundaries with the most used radiotracer [^18^F]fluorodeoxyglucose to obtain metabolic connectivity (M-MC). Twenty healthy participants underwent ultra-high-sensitivity fPET acquisitions on a large axial field-of-view scanner. M-MC was estimated using CompCor, spatio-temporal filtering, third-order polynomial detrending, baseline normalization and Euclidean distance at multiple temporal resolutions. mCov was assessed from SUVR images with subsequent network- or subject-specific matrix computation using independent component analysis, principal component analysis, or jackknife perturbation. Results demonstrate that while all MC methods are valid, CompCor and Euclidean distance perform optimally at high temporal resolutions (1-16s), whereas polynomial and spatio-temporal filters are more robust at lower sampling rates (>16s). mCov offers a population-level characterization of metabolic organization with the option to derive relative-to-group single-subject maps. This comparison provides methodological clarity and supports standardized use of molecular network analyses now integrated into the open-source fPET toolbox.

## Introduction

The human brain is organized into large-scale networks that support cognition, emotion, and behavior through dynamic interactions across distributed regions. Functional connectivity, typically assessed using functional magnetic resonance imaging (fMRI) or electroencephalography, has provided a framework for assessing such interactions by examining temporal correlations in spontaneous brain activity ^1, 2^. In particular, resting-state paradigms have revealed robust and reproducible network architectures, offering critical insights into both neuroscience and clinical applications. These approaches, however, primarily capture hemodynamic or electrophysiological signatures of neural activity, and therefore only reflect a portion of brain function ^3–6^.

Positron emission tomography (PET) provides a complementary perspective by probing molecular mechanisms of brain function. In both research and clinical settings, [^18^F]fluorodeoxyglucose ([^18^F]FDG) remains the gold standard for measuring glucose metabolism rate, a direct indicator of energy demand ^7, 8^. Conventional applications have used static images averaged across tens of minutes, which have been used to compute molecular covariance (mCov) across groups of subjects ^9–11^. Such across-subject covariance approaches, first explored in the 1980s ^10^ and 1990s ^12, 13^, quantify interindividual variation in tracer uptake between brain regions and have historically served as a proxy for molecular connectivity (MC). While widely applied to diverse tracers and straightforward to implement, mCov methods inherently capture population-level associations rather than temporal dynamics within individuals ^14^.

The last decade has seen advances in PET methodology and technology, enabling the transition from static to dynamic assessments of MC ^15–17^. The application of (bolus) + continuous infusion protocols combined with improved reconstruction algorithms and motion correction now allow functional PET (fPET) acquisitions with temporal resolutions on the order of seconds ^18, 19^. These developments, further accelerated by the introduction of large axial field-of-view (LAFOV) scanners with markedly increased sensitivity, provide the means to investigate spontaneous fluctuations in glucose metabolism akin to fMRI functional connectivity (FC) ^20^. This has established the foundation for estimating MC, which characterizes within-subject temporal associations of tracer uptake between regions during rest or task conditions. Such an approach holds potential to complement fMRI by offering a molecular-level counterpart of FC.

Alongside these advances, methodological diversity has expanded considerably. Multiple strategies for deriving metabolic connectivity (M-MC) from fPET data have focused on removing the cumulative component of the [^18^F]FDG signal, including spatio-temporal filters ^21, 22^, component-based noise correction ^19^, polynomial detrending ^17^, and normalization approaches ^15^. Other metrics exploit the unique kinetic properties by assessing the similarity of tracer time-activity curves, such as Euclidean distance ^23^. In parallel, mCov-based approaches remain valuable, as computation from static images not only offer group-level network characterizations but can also be leveraged to generate relative-to-group individual mCov matrices using methods such as independent component analysis (ICA) ^24^, principle component analysis (PCA) ^25^ or jackknife leave-one-out analysis ^26^. Although both approaches ultimately aim to estimate either individual-level MC, group-level mCov or relative individual mCov matrices, the choice of processing method substantially influences the resulting network patterns, their absolute magnitude, and their dependence on temporal resolutions or group composition, thus also determining the interpretation.

The present study provides a systematic comparison of widely used MC and mCov approaches exemplarily based on [^18^F]FDG as the most frequently used PET tracer in this neuroimaging. We assess their methodological assumptions, input requirements, and resulting output metrics across different temporal resolutions, to clarify their similarities and distinctions. This structured comparison is intended to guide interpretation, improve reproducibility, and define the conceptual boundaries of PET-based molecular network analyses. We have incorporated all of these methods into the fPET toolbox ^27^ to allow for a more unified and widespread use of these techniques.

## Materials and Methods

### Participants

Twenty healthy participants (mean age ± SD = 24.8 ± 3.6 years; 11 female) underwent a single [^18^F]FDG PET/CT scan using a next generation LAFOV scanner (Siemens Vision Quadra) followed by an MRI scan on a Siemens MAGNETOM Prisma 3T.

General health was evaluated through a structured medical assessment. This included medical history, physical examination, electrocardiogram, and routine laboratory testing. Psychiatric screening was conducted using the Structured Clinical Interview for DSM-IV Axis I Disorders to rule out current or past psychiatric diagnoses.

Exclusion criteria were any current or previous severe medical condition, psychiatric disorder, psychopharmacological treatment (past 6 months), or contraindications to PET or MRI. These included metallic implants, claustrophobia, and prior radiation exposure. Urine drug tests were performed at screening and before each scan. Additionally, female participants underwent urine pregnancy tests.

All participants provided written informed consent and received financial compensation for their time. The study was approved by the ethics committee of the Medical University of Vienna (EK1642/2022) and conducted in accordance with the Declaration of Helsinki.

### Study Design and data acquisition

For the LAFOV PET/CT study, participants were positioned with the head centered in the scanner bore to allow for ultra-high sensitivity acquisition (∼176 kcps/MBq) ^28, 29^. Each session began with a topogram and an ultra-low-dose CT scan (120 kVp, 20 mA, CareDose4D and CarekV enabled) being acquired after breath-hold instructions. CT images were reconstructed with 0.98 x 0.98 x 4 mm voxels ^30^. Thereafter, a 25-minute resting-state fPET acquisition was acquired in list-mode. During the scan, participants were instructed to remain awake with their eyes open, maintain fixation on a black crosshair presented on a gray background, and allow their mind to wander freely without focusing on any specific t. Radiotracer administration followed a bolus plus constant infusion protocol (5.1 MBq/kg body weight at a ratio of 20:80)^31^, initiated simultaneously with the start of PET acquisition.

T1-weighted images were recorded on a Siemens Prisma 3T scanner with a 64-channel head coil using the following parameters: TE/TR = 2.95/2300 ms, TI = 900 ms, flip angle = 9°, GRAPPA 2, 240 x 256 mm field of view, 176 slices, 1.05 x 1.05 x 1.20 mm3, TA = 5:09 min.

### Reconstruction and Preprocessing

The fPET data were reconstructed using Siemens’ e7 Tools. Data was reconstructed using 3D-TOF OP-OSEM with 4 iterations and 5 subsets into 1500 frames of 1s. This data was also rebinned into 16s and 30s frames, representing typical reconstructions performed in fPET research^16, 17, 22, 32^.

Preprocessing and quantification of all fPET data followed procedures established in previous studies ^31, 33^. In short, head motion correction was performed using SPM12 (https://www.fil.ion.ucl.ac.uk/spm) with the quality setting at “best”. Images were realigned to a motion-free reference created from a stable time segment late in the scan. The mean PET image was coregistered to each participant’s structural MRI, which in turn was spatially normalized to MNI space. These transformations were then applied to the full dynamic fPET series. Normalized data were spatially smoothed with an 8 mm Gaussian kernel.

### Parcellation and regional signal extraction

For both M-MC and mCov analyses, matrices were calculated from either the processed time series or standardized uptake value ratios (SUVR) images derived from each participant. SUVR images were created from the final 10 minutes of PET acquisition. Regional mean signals were extracted using a combined parcellation consisting of the Schaefer 100 cortical parcels ^34^ and 14 subcortical regions from the Harvard–Oxford atlas ^35^. The time series were truncated to exclude the initial 7.5 minutes and the final 1.5 minutes in order to eliminate the effects of the tracer bolus injection phase and any potential reconstruction artifacts at the end of acquisition.

### Molecular connectivity preprocessing

M-MC is derived at the within-subject level from dynamic fPET data by correlating regional time-activity curves (TACs), analogous to FC in fMRI. However, since the [^18^F]FDG signal accumulates over time, PET-specific preprocessing is necessary to account for low-frequency trends from cumulative tracer uptake. Depending on the approach, this either involves removing the uptake component to isolate spontaneous fluctuations or directly exploiting cumulative signal properties. Below we provide a brief description of the most commonly applied methods to compute metabolic connectivity. Table 1 provides a detailed overview of all M-MC and mCov methods tested, comparing inputs, outputs, and recommendations for function parametrization.

**Table 1:** Overview of all molecular connectivity (MC) and metabolic covariance (mCov) methods compared in the present study. Each approach differs in preprocessing complexity, and interpretational scope. MC methods estimate within-subject temporal associations from dynamic fPET data, whereas mCov methods quantify across-subject covariance from static PET images. Parameter selection and adaptation to temporal resolution are critical for obtaining valid and interpretable connectivity estimates.

| Method | Category | Output | Parameters | Advantages | Limitations | Optimal application and recommendation |
| --- | --- | --- | --- | --- | --- | --- |
| **Euclidean Distance** | MC | Distance/similarity matrix (1 – normalized Euclidean distance) |  | Simple and preprocessing-free; applicable across all temporal scales; robust to motion artifacts | Retains cumulative baseline uptake, which may bias connectivity magnitude; different interpretational basis than correlation methods | Feasible if high temporal resolution for correlation-based approaches is not available. |
| **CompCor** | MC | Residual time series → molecular connectivity matrix | Optional band-pass filter | Data-driven and flexible; removes structured noise (WM, CSF, motion) and low-frequency drift; preserves short-term metabolic fluctuations | May remove physiological signal at low temporal resolution if band-pass filtering is applied | Optimal at high temporal resolution ($\leq 16$ s); for coarser data, omit band-pass filtering |
| **Spatio-temporal Filter** | MC | Filtered TACs → molecular connectivity matrix | Temporal $\sigma_t$ and spatial $\sigma_s$ kernel widths | Dual-domain Gaussian smoothing; balances noise reduction and trend removal | Parameter-sensitive; incomplete baseline removal if kernel not tuned to frame duration | Works best at middle to low temporal resolution ($> 16$ s). Adapt temporal kernel for low resolutions ($\geq 30$ s) to avoid residual baseline components |
| **Third-order Polynomial** | MC | Detrended residuals → molecular connectivity matrix | Polynomial order | Simple and fast; widely implemented | Cannot model complex slow oscillations; edge artifacts at scan onset; reduced accuracy at extreme resolutions | Works best at middle to low temporal resolution ($> 16$ s). |
| **Baseline Normalization** | MC | Normalized residuals<br>→ molecular connectivity matrix |  | Removes global baseline drift; computationally minimal | Eliminates signal variance necessary for MC estimation; produces flat or unstructured matrices | Useful for baseline correction; not recommended for MC estimation |
| **Covariance** | mCov | Group-level covariance matrix |  | Simple and robust; tracer-independent; suitable for large samples | Lacks within-subject or temporal information; only population-level | Stable across tracers; ensure sufficient group size and biological specificity |
| **Independent Component Analysis (ICA)** | mCov | Independent spatial components; subject loadings | Number of components | Identifies independent metabolic networks; enables subject-level network expression maps | Component interpretation depends on decomposition stability and component number | Use fewer components than subjects; verify biological plausibility of components |
| **Principal Component Analysis (PCA)** | mCov | Principal component maps; subject scores; explained variance | Number of retained components | Reveals orthogonal axes of metabolic variance; interpretable variance structure | Orthogonality may separate overlapping or interacting networks | Select components based on explained variance |
| **Jackknife (Leave-one-out)** | mCov | Relative subject-specific covariance matrices; influence scores |  | Quantifies individual influence on group covariance; tests network stability and outlier detection | Computationally intensive for large N; sensitive to sample heterogeneity | Useful for outlier detection and patient characterization |

***Euclidean Distance*** ^23, 36^: Connectivity is computed as the similarity between TACs from different regions, defined as one minus the normalized Euclidean distance between TACs. Due to the heavy-tailed distribution of Euclidean similarity values, a Fisher z-transformation is applied to normalize the data. The values are then rescaled to the range [0, 1] to maintain consistency with the original scale ^23^. Unlike correlation-based methods, this approach does not require explicit baseline removal. Input is raw or preprocessed TACs, and the output is a similarity matrix constrained to positive values, reflecting molecular proximity across regions.

***CompCor*** ^19^: a component-based noise correction method adapted from fMRI ^37^. CompCor models structured variance from white matter, cerebrospinal fluid, and head motion, which are treated as nuisance regressors. Principal component analysis is used to extract dominant noise components from tissue-specific masks, which are then regressed out together with motion parameters and an optional temporal band-pass filter in a single step ^19, 38^. This combined approach effectively suppresses physiological and motion-related noise while simultaneously correcting for the cumulative tracer uptake. The output is a residual time series that retains moment-to-moment metabolic fluctuations relevant for connectivity estimation. For the present analysis, nuisance signals outside the 0.01-0.1 Hz frequency band were removed to isolate resting-state metabolic activity as based on previous work ^19^.

***Spatio-temporal filter*** ^22, 32^: applies Gaussian smoothing in both temporal and spatial dimensions to remove cumulative tracer uptake and low-frequency drift while retaining short-term signal fluctuations. Input consists of dynamic fPET data. The output is a filtered TAC set. The filter utilized a spatial Gaussian standard deviation of 1 voxel and a temporal Gaussian standard deviation of 2 frames as suggested previously ^22^.

***Third-order polynomial detrending*** ^17, 39^: regional TACs are fitted with a third-order polynomial function to model cumulative tracer uptake over time. The residuals derived from this polynomial fit represent the moment-to-moment signal fluctuations, facilitating subsequent calculations of inter-regional molecular connectivity.

***Baseline normalization*** ^15^: each TAC is normalized relative to the global brain signal at each time point, reducing interregional differences driven by baseline tracer uptake. Raw regional TACs are used as input.

### Molecular connectivity

For the Euclidian distance-based molecular connectivity (eMC) approach, no further calculation is required as the resulting similarity matrix already represents connectivity estimates. For the other approaches (CompCor, spatio-temporal filter, polynomial detrending, baseline normalization), connectivity was assessed by computing Pearson’s partial correlations between regional time courses (cMC), quantifying the linear association between each pair of brain regions while controlling for confounding signals. Here, head motion was accounted for by including the six realignment parameters as nuisance regressors in the partial correlation regression model (CompCor, see above) ^17, 19^ or in the partial correlations (polynomial detrending, spatio-temporal filter, and baseline normalization). This results in a connectivity matrix for each individual.

### Molecular Covariance

mCov is estimated at the between-subject level from PET data, reflecting interindividual variation in regional tracer uptake. Unlike MC, which requires dynamic acquisitions and preprocessing to address cumulative uptake, covariance computation operates on static images that e.g., summarize overall metabolism across the scan.

### Molecular Covariance Postprocessing

mCov provides a population-level view of network organization and can be further decomposed into relative, subject-specific or network-specific matrices using multivariate techniques.

***ICA*** ^24^: group-level ICA is applied to covariance matrices to extract independent spatial components reflecting metabolic networks. Input is region data for each subject, and the output is a set of independent metabolic networks with subject-specific loadings.

***PCA*** ^25^: reduces dimensionality by decomposing interregional covariance across subjects into orthogonal components. Input is the subject-by-region data matrix, and the output consists of principal components capturing dominant axes of metabolic variance, which can be interpreted as large-scale metabolic networks or each subject’s contribution to each component. Furthermore, the variance explained by each component can also be assessed.

***Leave-one-out jackknife*** ^26^: To assess robustness of covariance-derived networks, covariance or decomposition analyses are repeated iteratively while excluding one subject at a time. Inputs are static PET images across all subjects, and the output is a set of jackknife-derived covariance or component solutions. This enables the evaluation of stability of estimated networks as well as relative subject-specific metabolic covariance patterns.

### Statistics

To summarize findings across participants, individual correlation coefficients were Fisher z-transformed, averaged across subjects, and then back-transformed to obtain correlation values for group-level visualization. The resulting group-average connectivity and covariance matrices were compared using hierarchical clustering to evaluate similarities and differences between methods. Clustering employed Chebychev (maximum) distance as the dissimilarity metric, enabling visualization of the relative proximity between methods and quantitative assessment of their distinct connectivity patterns. Similarity between temporal resolutions and preprocessing methods was quantified by computing the correlation of the upper triangular elements of each method’s connectivity matrix. Eigenvalue decomposition was conducted to better assess the spatial structure of each method’s correlation/covariance matrix at each temporal resolution, and the results were visualized using a scree plot.

## Results

### Signal Characteristics across Metabolic Connectivity Processing Methods

To assess the impact of preprocessing strategies on the fPET signal, representative TACs were extracted from the visual cortex (Yeo-7 network) ^40^. Figure 1 illustrates the effect of frame length and filtering method on signal characteristics. Increasing frame duration reduced apparent noise across all M-MC processing methods, reflecting the trade-off between temporal resolution and signal stability. Since the Euclidean distance approach computes similarity directly from regional TACs without prior preprocessing, the cumulative baseline uptake remained, demonstrating that this method inherently incorporates the monotonic tracer accumulation component into its similarity estimation. In contrast, the CompCor approach effectively removed both the baseline uptake and high-frequency noise, yielding stable residual fluctuations across temporal resolutions. The CompCor-processed signal maintained a consistent amplitude between the 1s and 16s frames. At a temporal resolution of 30s, CompCor performance declined as expected, as the sampling rate no longer supported estimation of fluctuations within the 0.01-0.1 Hz frequency band, resulting in insufficient variance to estimate connectivity. Both the spatio-temporal and third-order polynomial filters behaved as effective high-pass filters, successfully removing the cumulative uptake component while preserving spontaneous fluctuations. Nevertheless, both methods exhibited residual low-frequency oscillations, particularly at coarser temporal resolutions. A pronounced signal drop was observed in the initial frames for both methods at 16s and 30s bins, suggesting edge effects introduced by the detrending procedures. The spatio-temporal filter performed better than the polynomial detrending in the 1s data, as slow oscillations were largely absent. In contrast, the baseline normalization approach no longer succeeded in fully removing cumulative tracer uptake. While the global signal spike during bolus injection was followed by an apparently stable time course, closer inspection of the connectivity-relevant portion of the signal revealed a consistent downward trend. This residual uptake drift introduced low-frequency bias into the signal, compromising its suitability for accurate metabolic connectivity estimation.

**Figure 1:**
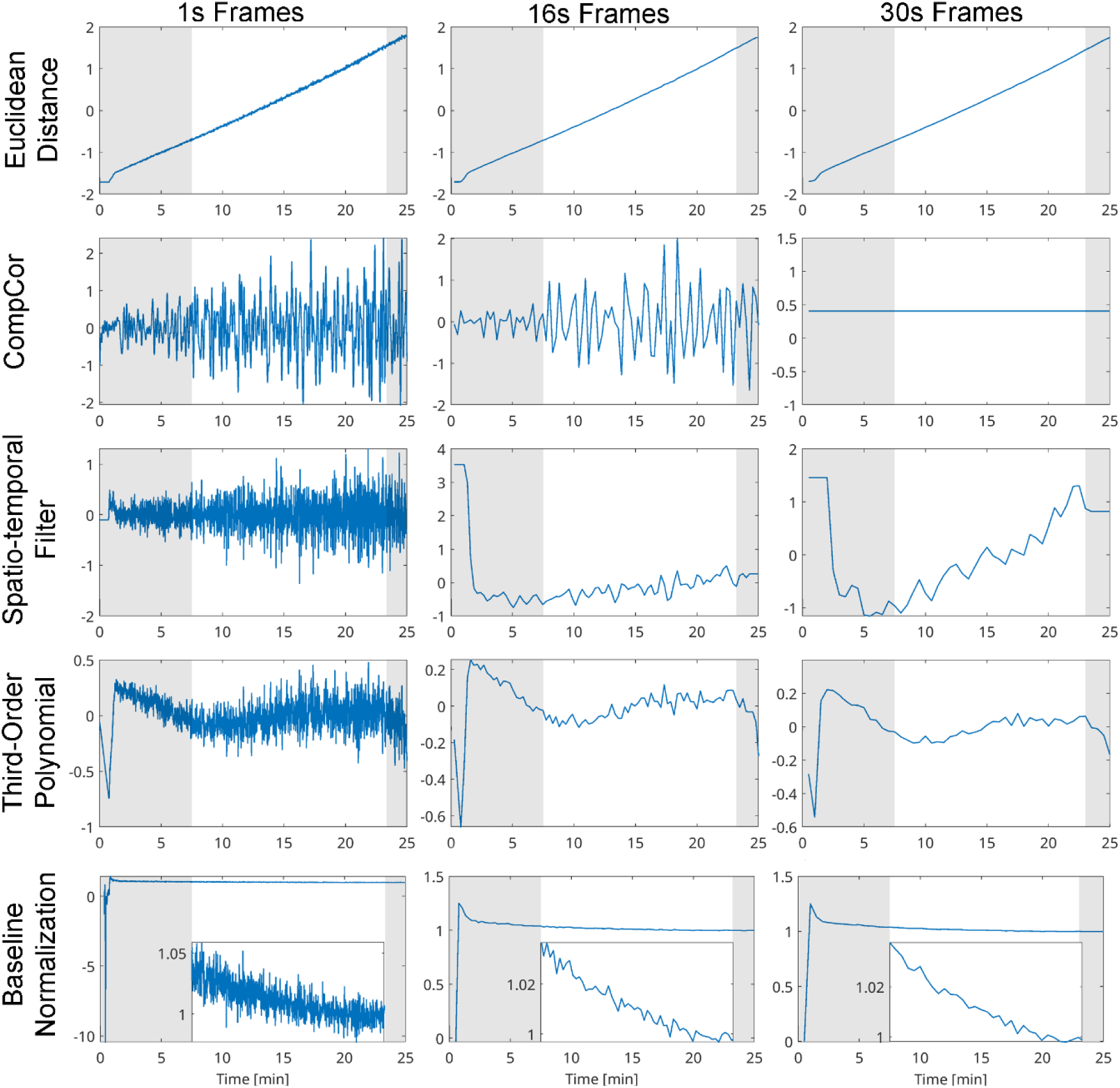
Representative time courses of metabolic connectivity preprocessing methods. [^18^F]FDG fPET signals from the visual cortex (Yeo-7 network) shown for 1 s, 16 s, and 30 s frame reconstructions. Each preprocessing technique demonstrates distinct handling of baseline uptake and temporal fluctuations. Euclidean distance metabolic connectivity, does not require baseline removal. CompCor effectively removes low-frequency drift and high-frequency noise, while polynomial and spatio-temporal filters behave as high-pass filters. Both the CompCor and spatio-temporal filters require parameter adjustments for optimal performance at lower temporal resolutions, exceeding 30s: CompCor processing benefits from excluding band-pass filtering during detrending, as an expected undersampling of 0.01-0.1 Hz fluctuations limited variance for estimating connectivity, whereas the spatio-temporal filter requires adaptation of the temporal kernel to the chosen frame duration. Baseline normalization fails to fully remove cumulative uptake: despite a large bolus-related spike at scan start, the subsequent signal shows a consistent downward trend, indicating residual tracer accumulation that biases connectivity estimation. The white area shows the signal used to estimate metabolic connectivity.

### Connectivity Patterns across Temporal Resolutions

The resulting connectivity matrices for each preprocessing method and temporal resolution are shown in Figure 2 and corresponding Eigenvalue decomposition in Supplementary Figure S1. Consistent with the signal characteristics described above, Euclidean distance produced similar connectivity patterns across all frame durations, with overall connectivity strength decreasing with lower temporal resolution. This pattern reflects reduced temporal variability and the influence of cumulative signal components at coarser binning. The CompCor method yielded structured network patterns for both the 1 s and 16 s data, showing enhanced contrast between major networks compared to other preprocessing strategies, see supplementary figure 1. Consistent with the degraded temporal frequency content observed in Figure 1, the 30s data did not retain usable connectivity structure within the 0.01-0.1 Hz range, resulting in near-zero and randomly structured correlations, due to the herein implemented bandpass limits. The third-order polynomial and spatio-temporal filters produced comparable connectivity matrices, exhibiting similar large-scale organization and interregional associations (Supplementary Table 1 and 2). However, for the 1s resolution, these approaches did not yield pronounced structural network organization. At the 30s temporal resolution, the third-order polynomial filter preserved slightly more structure and dynamic range than the spatio-temporal filter, suggesting that polynomial detrending may be less sensitive to low temporal sampling. The baseline normalization method produced connectivity matrices with the lowest network organization and connectivity amplitudes for all temporal resolutions. This appeared to be driven by residual tracer uptake rather than genuine metabolic fluctuations, reflecting that baseline normalization failed to completely remove cumulative signal components at lower sampling rates.

**Figure 2:**
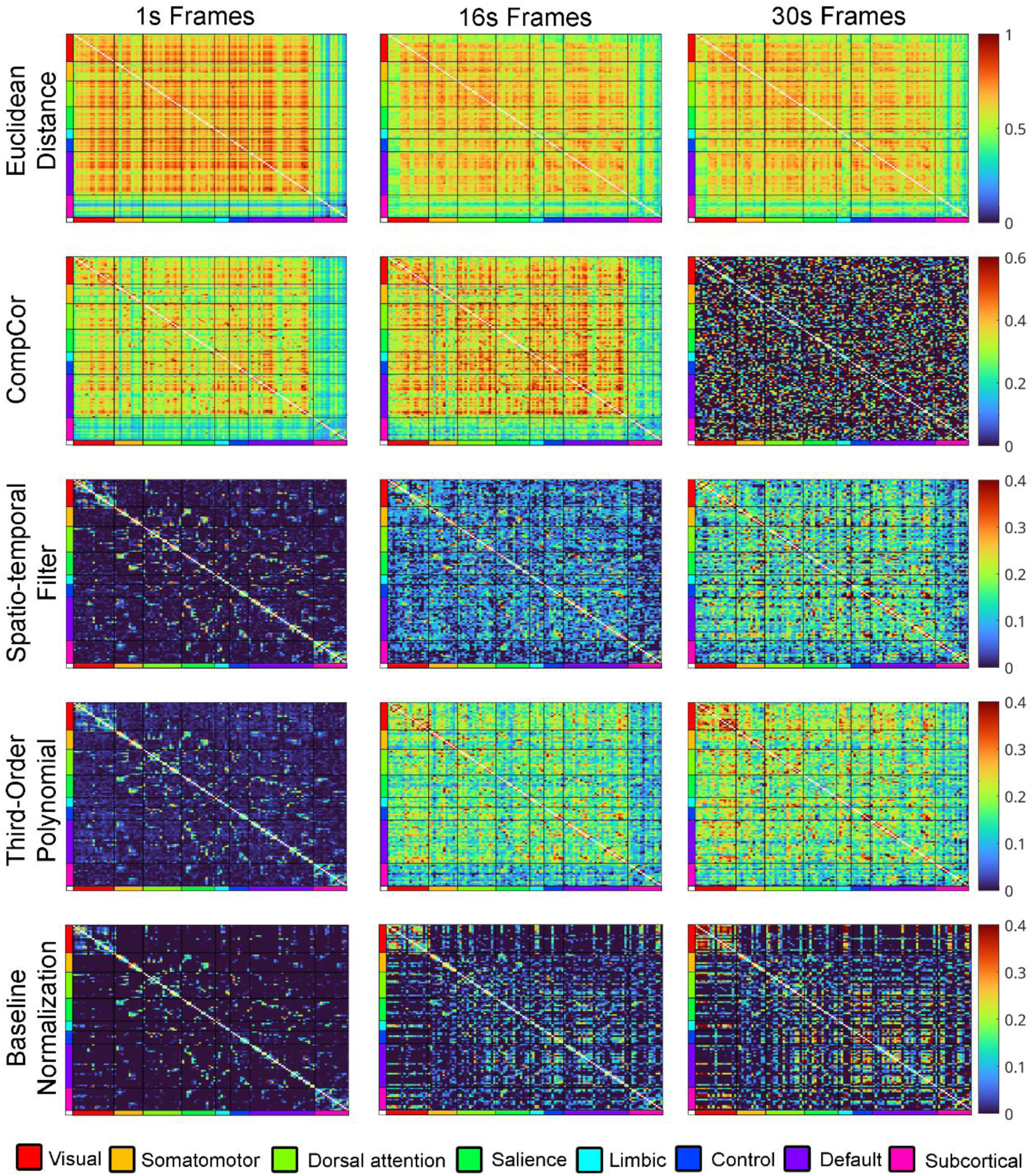
Metabolic connectivity matrices across temporal resolutions. Group-averaged connectivity matrices for each preprocessing approach. Euclidean distance produces consistent but baseline-influenced patterns across temporal resolutions. CompCor yields the most differentiated network structure at 1s and 16s, while polynomial and spatio-temporal filters produce similar connectivity profiles at 30s. Baseline normalization, however, produces patterns increasingly dominated by residual uptake, reflecting incomplete baseline removal rather than genuine connectivity. The CompCor with bandpass filtering yelds no discernable connectivity pattern at low resolutions (>30s), thus, removal or amending the filter cutoff frequencies would be necessary for accurate estimation.

### Metabolic Covariance Analysis

Figure 3a illustrates the group-level mCov matrix computed from static SUVR data, which can be further decomposed using several complementary analytical frameworks. ICA enables the extraction of distinct spatially independent subnetworks (Figure 3c) and subject-level component loadings representing individual network expression (Figure 3b). PCA similarly identifies spatial metabolic covariance patterns within the mCov matrix, revealing large-scale networks as principal components (Figure 3e). Each participant’s expression of these components can be quantified as a low-dimensional individual network contribution (Figure 3d), and the cumulative variance explained by successive components provides a measure of network dimensionality (Figure 3f). Perturbation of the mCov matrix through a leave-one-out jackknife procedure (Figure 3g–i) allows the derivation of relative subject-specific covariance maps and the identification of connections exhibiting high intersubject heterogeneity. This analysis also quantifies the contribution of each individual to the overall group covariance (Figure 3i), thereby characterizing the stability and subject-dependence of network topology.

**Figure 3:**
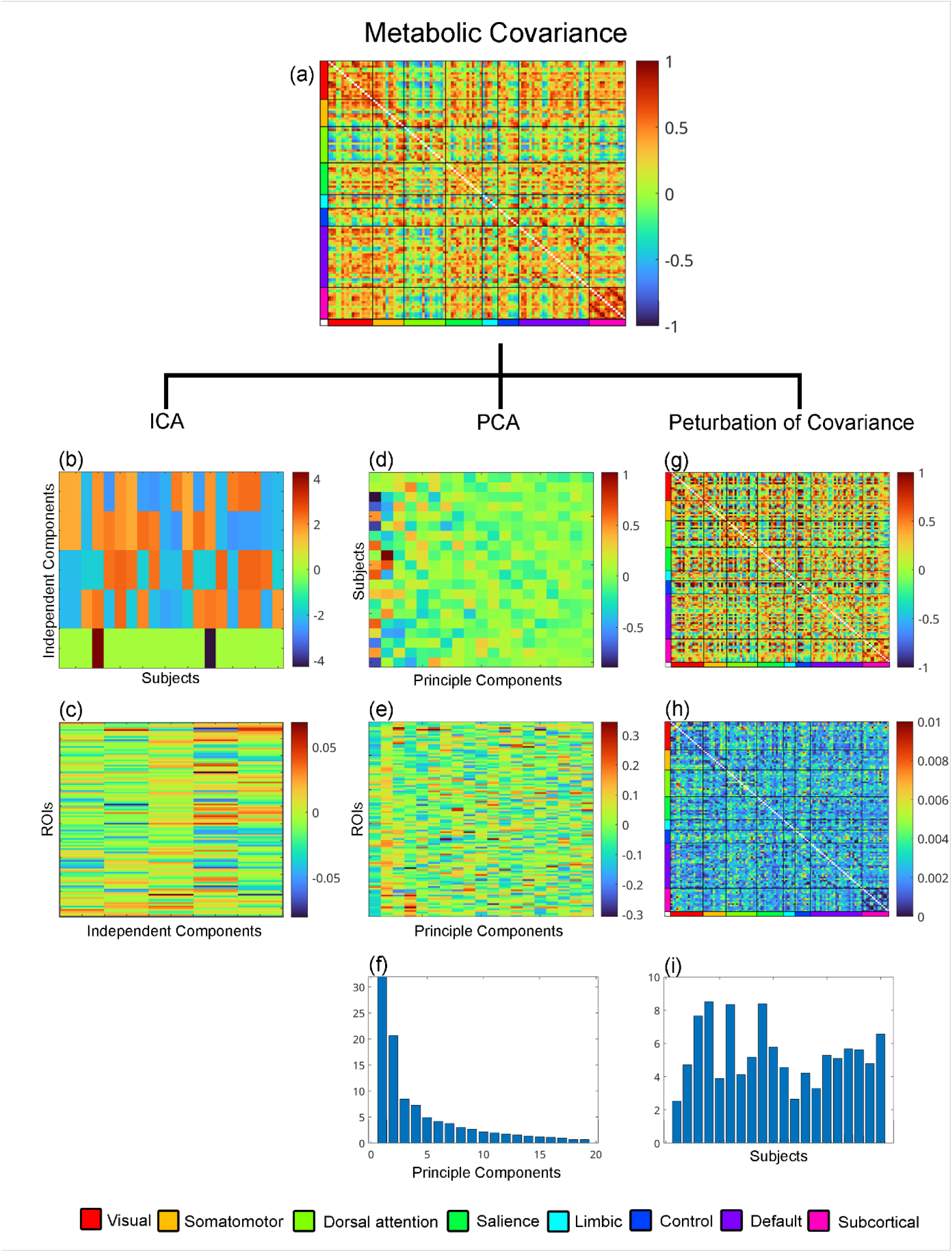
Metabolic covariance and decomposition approaches. (a) Group-level mCov matrix derived from static SUVR data. (b–c) Independent component analysis identifies spatially independent subnetworks and individual network expressions, in this case 5 independent components were chosen. (d–f) Principal component analysis reveals spatial covariance patterns, subject component loadings, and explained variance. (g–i) Leave-one-out jackknife analysis illustrates an exemplar subject-specific covariance patterns, and individual influence on group covariance.

### Clustering of Connectivity and Covariance Methods

Hierarchical clustering was used to compare the similarity of group-averaged connectivity and covariance matrices across all methods (Figure 4). The resulting linkage tree clearly separated the across-subject covariance-based approach from within-subject connectivity methods, highlighting their methodological and interpretational divergence. Among the M-MC techniques, the polynomial detrending and spatio-temporal filters clustered closely together, reflecting their similar temporal filtering characteristics. CompCor and Euclidean distance formed distinct branches, consistent with their differing treatment of baseline and noise components. Baseline normalization was positioned furthest from all other M-MC methods, indicating its distinct normalization properties and reduced network structure. Supplementary table 1 shows a numerical comparison of similarity between approaches.

**Figure 4:**
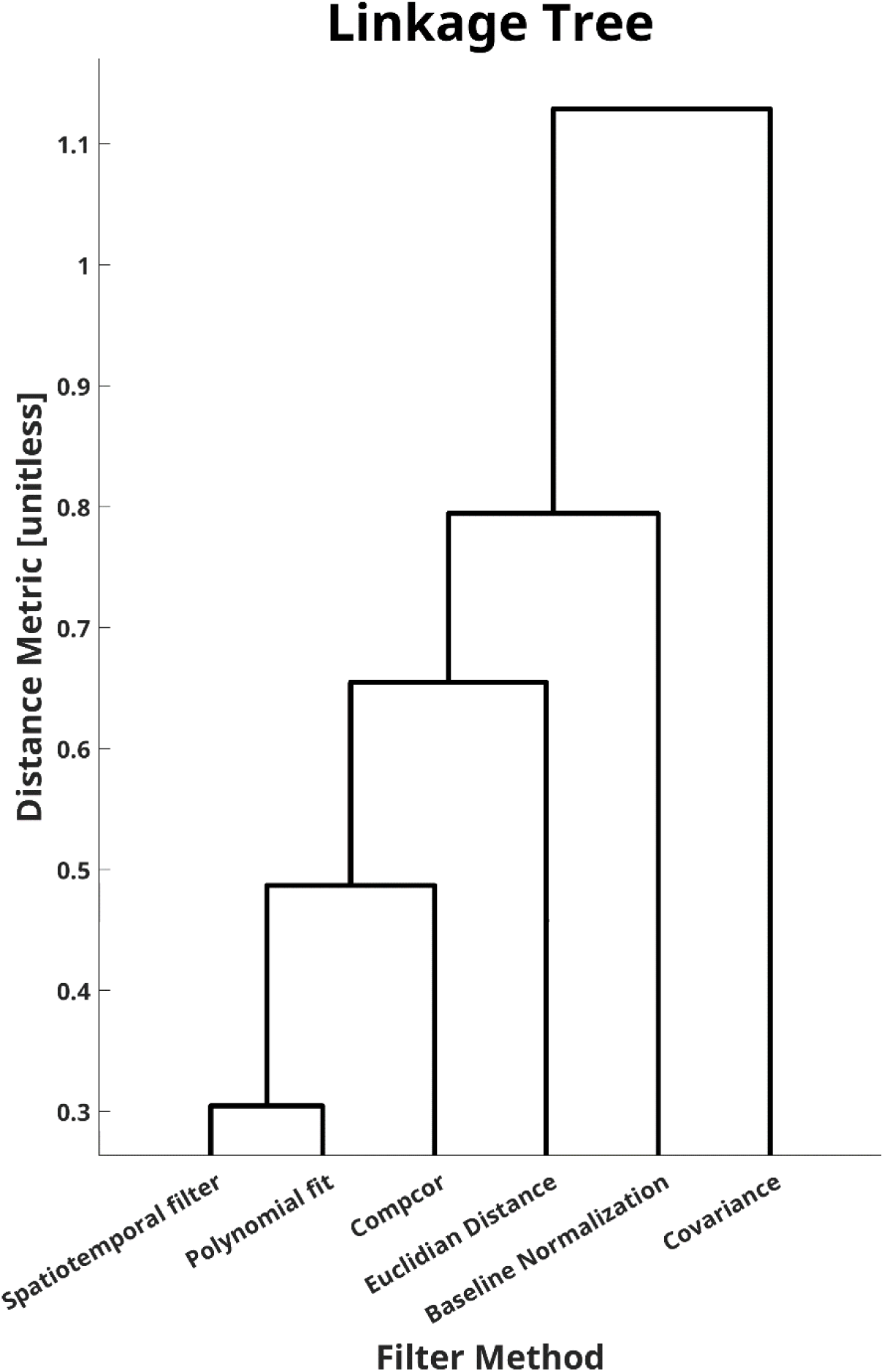
Hierarchical clustering of metabolic connectivity and covariance methods at a temporal resolution of 16s. Linkage tree depicting similarity across all methods using Chebychev distance. The dendrogram separates across-subject covariance approaches from within-subject connectivity methods. Among metabolic connectivity methods, polynomial and spatio-temporal filters cluster closely, CompCor and Euclidean distance form intermediate branches, and baseline normalization separates most distinctly.

## Discussion

Recent advances in high-temporal-resolution fPET have renewed interest in quantifying molecular network organization. However, the terminology surrounding MC and mCov has often been used interchangeably in the literature, leading to conceptual ambiguity ^16, 41, 42^. As emphasized in a recent consensus across numerous leading PET centers (2025) ^14^, the nomenclature has now been harmonized: MC refers to within-subject temporal associations derived from dynamic fPET data, whereas mCov describes across-subject associations derived from static or summary-level PET images. The present study continues these efforts by providing a thorough comparison of these two frameworks from a data-driven point of view. Using a dataset acquired at ultra-high sensitivity five MC and three mCov approaches were implemented to delineate their methodological differences and interpretational boundaries.

The comparison across M-MC preprocessing strategies highlights that most tested methods yield valid but method-specific representations of metabolic connectivity in humans, each with advantages for different temporal characteristics of the data. The Euclidean distance approach performed similar throughout all temporal resolutions. The method captures interregional similarity based on the overall shape of the TACs without requiring explicit detrending. However, because cumulative baseline uptake remains present, the resulting similarity values may be partially driven by shared global signal components, potentially inflating connectivity magnitudes. CompCor performed best at higher temporal resolutions (1s and 16s), capturing fine-grained temporal dynamics with reduced contamination from baseline uptake or noise. CompCor, by incorporating noise regression and band-pass filtering in a single model, efficiently removed low-frequency trends and high-frequency artifacts, producing stable and physiologically plausible fluctuations and structured network organization. As expected, at resolutions around 30s and lower, the often used frequency band of 0.01-0.1 does not yield feasible signal characteristics and connectivity patterns anymore. This issue can be easily resolved by setting the cutoff frequencies according to the study requirements. In contrast, polynomial detrending and spatio-temporal filtering performed well at intermediate temporal resolution (16s), yielding stable connectivity patterns with minimal baseline contamination. However, limitations emerged at both ends of the temporal spectrum. At high temporal resolution (1s), neither method sufficiently suppressed high-frequency noise, which propagated into the lack of network structure in M-MC estimates. At low temporal resolution (30s), the spatio-temporal filter failed to fully remove the cumulative baseline uptake, an effect that could likely be mitigated by adapting the spatial and/or temporal kernel parameters. The polynomial detrending approach, in turn, was unable to resolve slower oscillatory components across all temporal resolutions, thereby compromising the accurate assessment of M-MC. This observed limitation is consistent with previous reports showing that low-order polynomial filtering may be insufficient to model the complex temporal dynamics of fPET signals at resting state^43^, therefore recommending alternative preprocessing strategies for the computation of connectivity. The baseline normalization approach, was found to be insufficient for isolating metabolic fluctuations in the current data. Although the method reduced the overall amplitude of tracer uptake, it retained a consistent upward or downward drift, depending on region, that propagated into the connectivity estimation, producing network patterns driven by these effects rather than the spontaneous fluctuations in glucose metabolism. This becomes apparent particularly at lower temporal resolutions.

Overall, each preprocessing approach entails distinct advantages and limitations depending on the temporal characteristics of the acquisition. Techniques such as CompCor and Euclidean distance are particularly suited for fPET data with high(er) temporal framing. The Euclidean distance method offers a simple, “processing-free” alternative that performs consistently across temporal resolutions. Because it quantifies the distance between regional TACs rather than their temporal correlation, it provides a fundamentally different perspective on interregional similarity that can capture broader kinetic relationships. Nevertheless, the interpretation of its resulting connectivity metrics differs from correlation-based approaches, as cumulative baseline components remain embedded in the similarity estimates. CompCor performs best when sufficient temporal resolution allows for data-driven extraction of structured noise components. It flexibly adapts to the frequency range of interest and efficiently removes low- and high-frequency noise within a single regression model. For lower temporal resolutions, however, the inclusion of the optional band-pass filter can excessively remove the signal, and regressing out only the nuisance tissue components may be more appropriate. The polynomial detrending approach is a simpler alternative, albeit with limited flexibility in modelling signal trends and may fail to remove slow oscillatory dynamics in the fPET signal. In contrast, the spatio-temporal filter allows for parameterized control of temporal and spatial kernels, offering better adaptability to data acquired at lower temporal resolutions. Adjusting these kernel parameters to the frame duration is essential to ensure adequate baseline removal without removing physiologically relevant fluctuations. The choice of preprocessing method should therefore be guided by both the acquisition design and the expected frequency content of metabolic fluctuations. High-temporal fPET acquisitions benefit from adaptive, data-driven filtering approaches, whereas simpler polynomial or kernel-based filters may be sufficient when sampling rates are lower or computational simplicity is prioritized.

mCov analyses offer a conceptually distinct yet complementary framework. In contrast to MC, which relies on dynamic data, mCov requires only static PET images, such as SUVR, binding potential, or total distribution volume, and can therefore be applied to a broad range of radiotracers and study designs. This flexibility, alongside relatively low methodological demands, makes mCov particularly suitable for large-scale or multimodal datasets where dynamic acquisitions are impractical. Its simplicity also promotes reproducibility, as the covariance structure can be derived directly from existing static imaging cohorts without specialized acquisition protocols. Conceptually, mCov captures interindividual co-variation in regional tracer uptake, reflecting stable, trait-like patterns of the molecular network organization across a population. However, by design, it lacks a temporal dimension and therefore cannot probe within-subject dynamics or rapid fluctuations in metabolic coupling. Its interpretability is thus restricted to population-level associations i.e. identifying how regions covary across individuals rather than how they interact over time within an individual. Extensions of this framework aim to extract more detailed information from the group covariance structure. Decomposition methods such as ICA and PCA enable the identification of spatially independent or orthogonal metabolic networks, respectively, while quantifying each subject’s contribution to the network. ICA isolates non-overlapping subnetworks driven by shared interindividual variance, providing biologically interpretable motifs of metabolic organization. PCA, in turn, offers a dimensionality-reduced representation of the covariance space, highlighting the principal axes of variance that explain global and regional metabolic relationships. These decompositions transform static mCov data into interpretable network maps and facilitate cross-subject comparisons in health and disease. Perturbation-based analyses, including jackknife leave-one-out procedures, extend mCov’s interpretability further by estimating the stability and subject-specific influence on group-level covariance topology. By iteratively removing individual subjects and recalculating the covariance matrix, these methods reveal which connections are consistent or heterogeneous across participants, and which individuals disproportionately shape the group structure. Such robustness assessments are particularly valuable in heterogeneous clinical samples, where outlier contributions or subgroup effects may otherwise remain obscured. Still, it needs to be highlighted that individual mCov matrices obtained with a jackknife approach can only be interpreted in relation to the underlying group.

In summary, mCov operates from the group downward, defining a shared covariance structure and then assessing individual deviations relative to it, whereas MC operates from the individual upward, aggregating temporal associations across regions to infer group-level trends. These complementary inferential directions delineate the conceptual and methodological boundary between dynamic and static molecular network analyses, emphasizing that both perspectives are necessary to achieve a comprehensive understanding of large-scale metabolic organization in the human brain.

The empirical results from the present comparison reinforce this conceptual distinction. Hierarchical clustering revealed a clear separation between mCov-based and MC-based methods, mirroring their differing analytical hierarchies and data dependencies. Whereas mCov clustered as across-subject framework insensitive to temporal sampling, MC methods formed distinct branches reflecting their preprocessing strategies and frequency sensitivities. Notably, the close association between the polynomial and spatio-temporal filters underscores their similar design and outputs, while CompCor and Euclidean distance occupy intermediate positions. This pattern demonstrates that methodological choices, not only in preprocessing but also in the inferential framework, directly shape the resulting network architecture. Together, these findings highlight that mCov and MC are not interchangeable measures of “connectivity,” but complementary frameworks that capture different organizational dimensions of molecular brain networks.

Together, these findings provide a comprehensive comparison of molecular connectivity and covariance frameworks. MC approaches offer individual-level, temporally resolved assessments of metabolic interactions but require careful preprocessing tailored to temporal resolution. In contrast, mCov methods provide robust, population-level descriptions of molecular organization applicable across radiotracers but lack within-subject dynamics. Each method thus occupies a distinct position in the molecular connectomics landscape, with its own analytical strengths and constraints. Standardizing terminology and clarifying these methodological boundaries will enhance reproducibility and comparability across studies, facilitating the integration of molecular and functional imaging into a unified framework of brain network organization. To further enhance reproducibility and allow for cohesive analyses pipelines, each of the MC and mCov methods have been integrated into the fPET toolbox ^27^.

## Statements and Declarations

**Clinical trial number**: NCT06243783

### Author Contribution

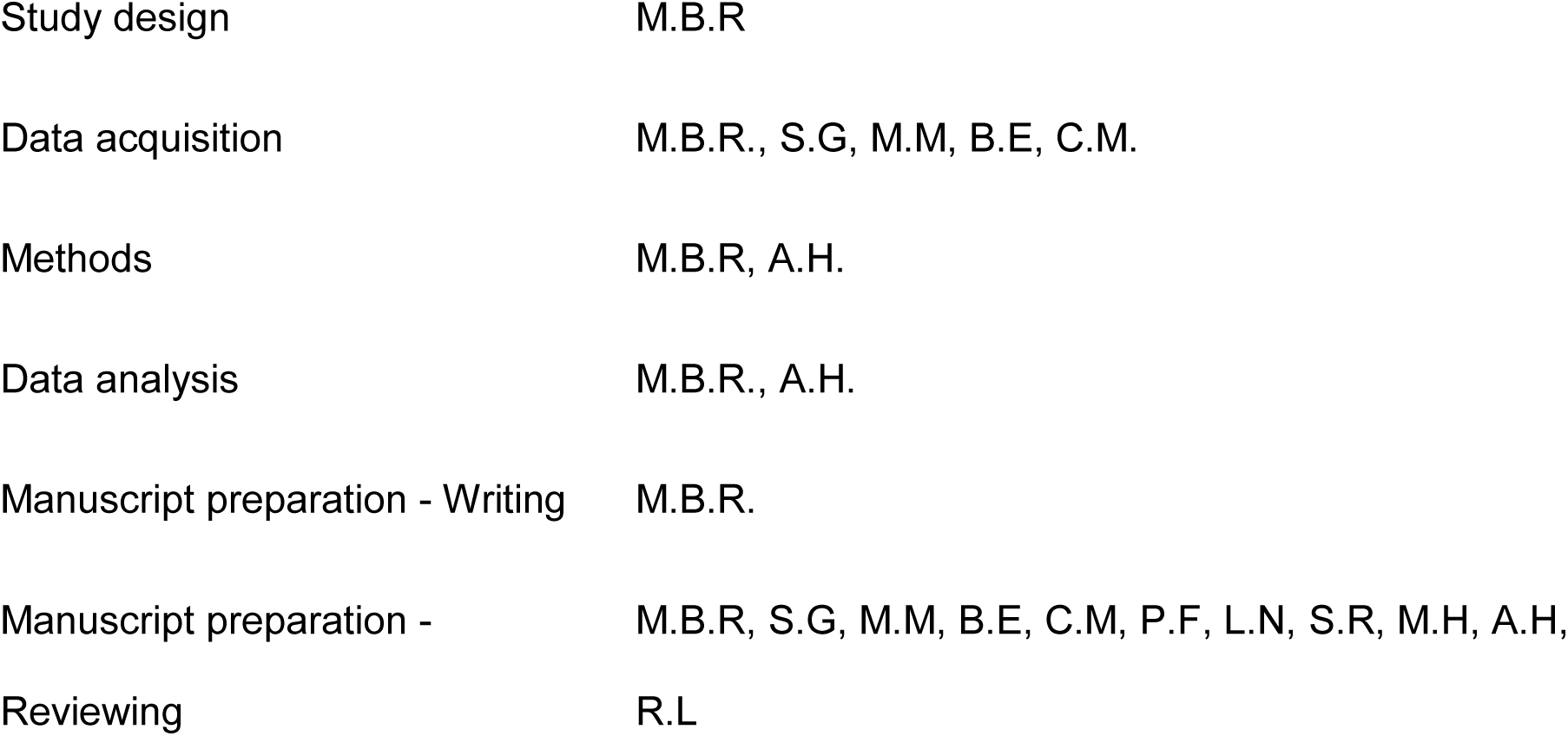

All authors discussed the implications of the findings and approved the final version of the manuscript.

### Ethics approval

This study was approved by the ethics committee of the Medical University of Vienna (EK 1642/2022) and conducted in accordance with the Declaration of Helsinki.

### Consent to participate

At the screening visit, all participants provided written informed consent after detailed explanation of the study protocol.

### Consent to publish

Not applicable.

### Funding

This research was funded in whole, or in part, by the Austrian Science Fund (FWF) [grant DOI: 10.55776/KLI1006 and 10.55776/PAT6608924, PI: R. Lanzenberger, and grant DOI: 10.55776/KLI1151 and 10.55776/PAT5436523, PI: A. Hahn]. Christian Milz is a recipient of a DOC Fellowship (27221) from the Austrian Academy of Sciences at the Department of Psychiatry and Psychotherapy, Medical University of Vienna. For open access purposes, the author has applied a CC BY public copyright license to any author accepted manuscript version arising from this submission.

### Disclosure / Conflict of Interest

R. Lanzenberger received investigator-initiated research funding from Siemens Healthcare regarding clinical research using PET/MR and travel grants and/or conference speaker honoraria from Janssen-Cilag Pharma GmbH in 2023, and Bruker BioSpin, Shire, AstraZeneca, Lundbeck A/S, Dr. Willmar Schwabe GmbH, Orphan Pharmaceuticals AG, Janssen-Cilag Pharma GmbH, Heel and Roche Austria GmbH., and Janssen-Cilag Pharma GmbH in the years before 2020. He is a shareholder of the start-up company BM Health GmbH, Austria since 2019. M. Hacker received consulting fees and/or honoraria from Bayer Healthcare BMS, Eli Lilly, EZAG, GE Healthcare, Ipsen, ITM, Janssen, Roche, and Siemens Healthineers. All other authors declare no potential conflicts of interest with respect to the research, authorship, and/or publication of this article.

### Data Availability Statement

Raw data will not be made publicly available due to reasons of data protection. Processed data can be obtained from the corresponding author with a data-sharing agreement, approved by the departments of legal affairs and data clearing of the Medical University of Vienna. Code is freely available within the fPET toolbox and can be downloaded from: https://github.com/NeuroimagingLabsMUV/fPET-toolbox

## References

1. Biswal B, Yetkin FZ, Haughton VM, et al. Functional connectivity in the motor cortex of resting human brain using echo-planar MRI. Magn Reson Med 1995; 34: 537–541. 1995/10/01. DOI: 10.1002/mrm.1910340409.

2. Reed MB, Klöbl M, Godbersen GM, et al. Serotonergic modulation of effective connectivity in an associative relearning network during task and rest. NeuroImage 2022; 249: 118887. DOI: 10.1016/j.neuroimage.2022.118887.

3. Buzsáki G, Anastassiou CA and Koch C. The origin of extracellular fields and currents — EEG, ECoG, LFP and spikes. Nature Reviews Neuroscience 2012; 13: 407–420. DOI: 10.1038/nrn3241.

4. Fox PT and Raichle ME. Focal physiological uncoupling of cerebral blood flow and oxidative metabolism during somatosensory stimulation in human subjects. Proc Natl Acad Sci U S A 1986; 83: 1140–1144. 1986/02/01. DOI: 10.1073/pnas.83.4.1140.

5. Raichle ME and Mintun MA. Brain work and brain imaging. Annu Rev Neurosci 2006; 29: 449–476. 2006/06/17. DOI: 10.1146/annurev.neuro.29.051605.112819.

6. Sundqvist N, Sten S, Thompson P, et al. Mechanistic model for human brain metabolism and its connection to the neurovascular coupling. PLoS Comput Biol 2022; 18: e1010798. 2022/12/23. DOI: 10.1371/journal.pcbi.1010798.

7. Sokoloff L. The physiological and biochemical bases of functional brain imaging. Cogn Neurodyn 2008; 2: 1–5. 2008/11/13. DOI: 10.1007/s11571-007-9033-x.

8. Vaishnavi SN, Vlassenko AG, Rundle MM, et al. Regional aerobic glycolysis in the human brain. Proc Natl Acad Sci U S A 2010; 107: 17757–17762. 2010/09/15. DOI: 10.1073/pnas.1010459107.

9. Hahn A, Haeusler D, Kraus C, et al. Attenuated serotonin transporter association between dorsal raphe and ventral striatum in major depression. Hum Brain Mapp 2014; 35: 3857–3866. 2014/01/21. DOI: 10.1002/hbm.22442.

10. Horwitz B, Duara R and Rapoport SI. Intercorrelations of glucose metabolic rates between brain regions: application to healthy males in a state of reduced sensory input. J Cereb Blood Flow Metab 1984; 4: 484–499. 1984/12/01. DOI: 10.1038/jcbfm.1984.73.

11. Yakushev I, Ripp I, Wang M, et al. Mapping covariance in brain FDG uptake to structural connectivity. European Journal of Nuclear Medicine and Molecular Imaging 2022; 49: 1288–1297. DOI: 10.1007/s00259-021-05590-y.

12. Horwitz B, McIntosh AR, Haxby JV, et al. Network analysis of brain cognitive function using metabolic and blood flow data. Behavioural Brain Research 1995; 66: 187–193. DOI: 10.1016/0166-4328(94)00139-7.

13. McIntosh AR, Grady CL, Ungerleider LG, et al. Network analysis of cortical visual pathways mapped with PET. J Neurosci 1994; 14: 655–666. 1994/02/01. DOI: 10.1523/jneurosci.14-02-00655.1994.

14. Reed MB, Cocchi L, Sander CY, et al. Connecting the dots: approaching a standardized nomenclature for molecular connectivity in positron emission tomography. European Journal of Nuclear Medicine and Molecular Imaging 2025. DOI: 10.1007/s00259-025-07357-1.

15. Ionescu TM, Amend M, Hafiz R, et al. Elucidating the complementarity of resting-state networks derived from dynamic [(18)F]FDG and hemodynamic fluctuations using simultaneous small-animal PET/MRI. Neuroimage 2021; 236: 118045. 2021/04/14. DOI: 10.1016/j.neuroimage.2021.118045.

16. Jamadar SD and Egan GF. Resting-State FDG-PET Connectivity: Covariance, Ergodicity, and Biomarkers. Response to Commentary by Sala et al.; Static versus Functional PET: Making Sense of Metabolic Connectivity. Cereb Cortex 2022; 32: 2054–2055. 2021/10/07. DOI: 10.1093/cercor/bhab316.

17. Reed MB, Ponce de León M, Vraka C, et al. Whole-body metabolic connectivity framework with functional PET. Neuroimage 2023; 271: 120030. 2023/03/17. DOI: 10.1016/j.neuroimage.2023.120030.

18. Hahn A, Reed MB, Vraka C, et al. High-temporal resolution functional PET/MRI reveals coupling between human metabolic and hemodynamic brain response. Eur J Nucl Med Mol Imaging 2024; 51: 1310–1322. 2023/12/06. DOI: 10.1007/s00259-023-06542-4.

19. Reed MB, Graf S, Murgaš M, et al. High-temporal resolution metabolic connectivity resolved by component-based noise correction. bioRxiv 2025: 2025.2008.2018.670788.DOI: 10.1101/2025.08.18.670788.

20. Hahn A, Falb P, Murgas M, et al. [18F]FDG Revisited: A New Perspective on the Temporal Dynamics of Brain Glucose Metabolism. Preprints. Preprints, 2025.

21. Jamadar SD, Ward PGD, Close TG, et al. Simultaneous BOLD-fMRI and constant infusion FDG-PET data of the resting human brain. Sci Data 2020; 7: 363. 2020/10/23. DOI: 10.1038/s41597-020-00699-5.

22. Jamadar SD, Ward PGD, Liang EX, et al. Metabolic and Hemodynamic Resting-State Connectivity of the Human Brain: A High-Temporal Resolution Simultaneous BOLD-fMRI and FDG-fPET Multimodality Study. Cereb Cortex 2021; 31: 2855–2867. 2021/02/03. DOI: 10.1093/cercor/bhaa393.

23. Volpi T, Vallini G, Silvestri E, et al. A new framework for metabolic connectivity mapping using bolus [(18)F]FDG PET and kinetic modeling. J Cereb Blood Flow Metab 2023; 43: 1905–1918. 2023/06/28. DOI: 10.1177/0271678x231184365.

24. Di X and Biswal BB. Metabolic brain covariant networks as revealed by FDG-PET with reference to resting-state fMRI networks. Brain Connect 2012; 2: 275–283. 2012/10/03. DOI: 10.1089/brain.2012.0086.

25. Spetsieris PG and Eidelberg D. Spectral guided sparse inverse covariance estimation of metabolic networks in Parkinson’s disease. Neuroimage 2021; 226: 117568. 2020/11/28. DOI: 10.1016/j.neuroimage.2020.117568.

26. Yao Z, Hu B, Chen X, et al. Learning Metabolic Brain Networks in MCI and AD by Robustness and Leave-One-Out Analysis: An FDG-PET Study. American Journal of Alzheimer’s Disease & Other Dementias® 2017; 33: 42–54. DOI: 10.1177/1533317517731535.

27. Hahn A, Reed MB, Milz C, et al. A unified approach for identifying PET-based neuronal activation and molecular connectivity with the functional PET toolbox. J Cereb Blood Flow Metab 2025: 271678x251370831. 2025/09/08. DOI: 10.1177/0271678x251370831.

28. Pommranz CM, Elmoujarkach EA, Lan W, et al. A digital twin of the Biograph Vision Quadra long axial field of view PET/CT: Monte Carlo simulation and image reconstruction framework. EJNMMI Physics 2025; 12: 31. DOI: 10.1186/s40658-025-00738-3.

29. Prenosil GA, Sari H, Fürstner M, et al. Performance Characteristics of the Biograph Vision Quadra PET/CT system with long axial field of view using the NEMA NU 2-2018 Standard. Journal of Nuclear Medicine 2021: jnumed.121.261972. DOI: 10.2967/jnumed.121.261972.

30. Reed MB, Godbersen GM, Vraka C, et al. Comparison of cardiac image-derived input functions for quantitative whole body [(18)F]FDG imaging with arterial blood sampling. Front Physiol 2023; 14: 1074052. 2023/04/11. DOI: 10.3389/fphys.2023.1074052.

31. Rischka L, Gryglewski G, Pfaff S, et al. Reduced task durations in functional PET imaging with [(18)F]FDG approaching that of functional MRI. Neuroimage 2018; 181: 323–330. 2018/07/04. DOI: 10.1016/j.neuroimage.2018.06.079.

32. Deery HA, Liang EX, Moran C, et al. Metabolic connectivity has greater predictive utility for age and cognition than functional connectivity. Brain Communications 2025; 7: fcaf075. DOI: 10.1093/braincomms/fcaf075.

33. Hahn A, Breakspear M, Rischka L, et al. Reconfiguration of functional brain networks and metabolic cost converge during task performance. eLife 2020; 9 2020/04/22. DOI: 10.7554/eLife.52443.

34. Schaefer A, Kong R, Gordon EM, et al. Local-Global Parcellation of the Human Cerebral Cortex from Intrinsic Functional Connectivity MRI. Cerebral Cortex 2018; 28: 3095–3114. DOI: 10.1093/cercor/bhx179.

35. Desikan RS, Ségonne F, Fischl B, et al. An automated labeling system for subdividing the human cerebral cortex on MRI scans into gyral based regions of interest. Neuroimage 2006; 31: 968–980. 2006/03/15. DOI: 10.1016/j.neuroimage.2006.01.021.

36. Vallini G, Silvestri E, Volpi T, et al. Individual-level metabolic connectivity from dynamic [(18)F]FDG PET reveals glioma-induced impairments in brain architecture and offers novel insights beyond the SUVR clinical standard. Eur J Nucl Med Mol Imaging 2025; 52: 836–850. 2024/10/30. DOI: 10.1007/s00259-024-06956-8.

37. Behzadi Y, Restom K, Liau J, et al. A component based noise correction method (CompCor) for BOLD and perfusion based fMRI. NeuroImage 2007; 37: 90–101. DOI: 10.1016/j.neuroimage.2007.04.042.

38. Klöbl M, Seiger R, Vanicek T, et al. Escitalopram modulates learning content-specific neuroplasticity of functional brain networks. NeuroImage 2022; 247: 118829. DOI: 10.1016/j.neuroimage.2021.118829.

39. Hahn A, Gryglewski G, Nics L, et al. Quantification of Task-Specific Glucose Metabolism with Constant Infusion of 18F-FDG. Journal of Nuclear Medicine 2016; 57: 1933. DOI: 10.2967/jnumed.116.176156.

40. Yeo BT, Krienen FM, Sepulcre J, et al. The organization of the human cerebral cortex estimated by intrinsic functional connectivity. J Neurophysiol 2011; 106: 1125–1165. 2011/06/10. DOI: 10.1152/jn.00338.2011.

41. Sala A, Lizarraga A, Ripp I, et al. Static versus Functional PET: Making Sense of Metabolic Connectivity. Cerebral Cortex 2022; 32: 1125–1129. DOI: 10.1093/cercor/bhab271.

42. Severino M, Peretti DE, Bardiau M, et al. Molecular connectivity studies in neurotransmission: a scoping review. Imaging Neurosci (Camb) 2025; 3 2025/08/13. DOI: 10.1162/imag_a_00530.

43. Coursey SE, Mandeville J, Reed MB, et al. On the analysis of functional PET (fPET)-FDG: Baseline mischaracterization can introduce artifactual metabolic (de)activations. Imaging Neuroscience 2025; 3: IMAG.a.110. DOI: 10.1162/IMAG.a.110.

