## Supplementary Figure 1 for "A Comparative Evaluation of Molecular Connectivity and Covariance Approaches"

### *Supplementary Material*

**Running Title: Metabolic connectivity and covariance approaches**

**# Correspondence to:**
Assoc.Prof. PD. Dr. Andreas Hahn, MSc

ORCID: https://orcid.org/0000-0001-9727-7580
Medical University of Vienna, Department of Psychiatry and Psychotherapy, Austria

| | Euclidean Distance | Compcor | Spatio temporal | Polynomial | Baseline Normalization | Covariance |
|---|---|---|---|---|---|---|
| **Euclidean Distance** | 1.000 | 0.652 | 0.365 | 0.503 | -0.010 | 0.199 |
| **Compcor** | | 1.000 | 0.447 | 0.726 | -0.005 | 0.026 |
| **Spatio temporal** | | | 1.000 | 0.733 | -0.001 | 0.181 |
| **Polynomial** | | | | 1.000 | -0.005 | 0.108 |
| **Baseline Normalization** | | | | | 1.000 | 0.009 |
| **Covariance** | | | | | | 1.000 |

Supplementary Table 1: Comparison of similarity between all metabolic connectivity and covariance matrices at 16s temporal resolution. This table reports the pairwise correlation coefficients among metabolic connectivity matrices derived using different filtering and preprocessing strategies (Spatiotemporal, Polynomial, Baseline Normalization, Euclidean Distance, CompCor, and Covariance). Values represent the degree of similarity between the resulting connectivity patterns across methods. Higher correlation values indicate greater concordance between methods in capturing underlying metabolic connectivity structure.

| Time Window Comparison | Spatiotemporal | Polynomial | Baseline Normalization | Euclidean Distance | Compcor |
|---|---|---|---|---|---|
| **1s vs 16s** | 0.68 | 0.54 | 0.66 | 0.80 | 0.88 |
| **1s vs 30s** | 0.54 | 0.51 | 0.58 | 0.81 | 0.00 |
| **16s vs 30s** | 0.58 | 0.67 | 0.88 | 0.97 | -0.01 |

Supplementary Table 2: Comparison of metabolic connectivity matrices between temporal resolutions within each of the approaches. This table presents the correlation coefficients between metabolic connectivity matrices computed at different temporal resolutions (1 s, 16 s, and 30 s) across five filtering strategies: Spatiotemporal, Polynomial, Baseline Normalization, Euclidean Distance, and CompCor. Each value represents the degree of similarity in the resulting connectivity patterns between pairs of temporal resolutions. Higher values indicate stronger correspondence in connectivity structure across time windows.

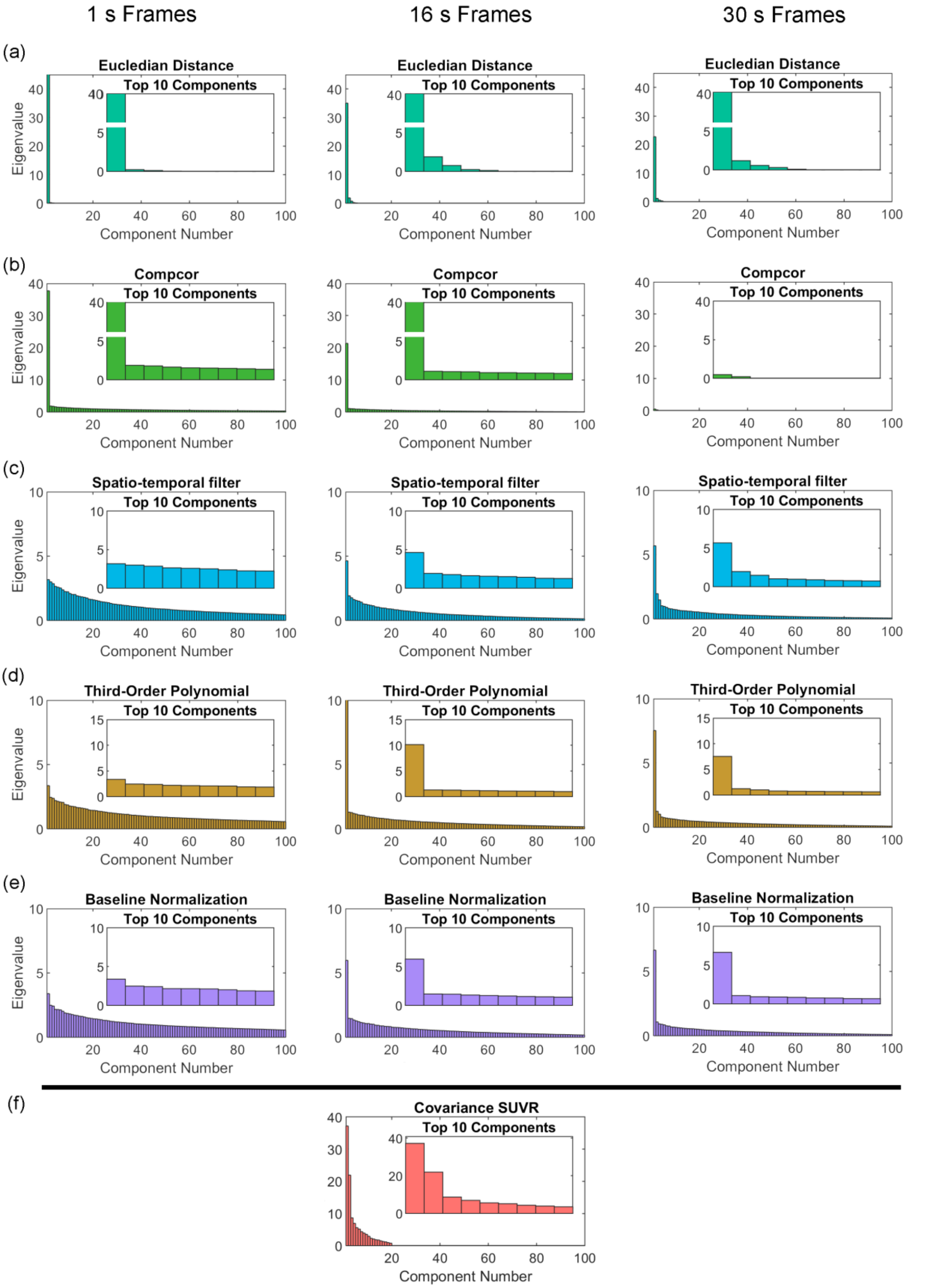


Supplementary Figure 1: Overview of eigenvalue decompositions across filtering methods and temporal resolutions. This figure illustrates the eigenvalue spectra derived from metabolic connectivity matrices processed using six different filtering and preprocessing strategies at varying temporal resolutions. A high eigenvalue component indicates the presence of dominant, structured patterns within the data. Prominent eigenvalue components are observed for (a) the Euclidean Distance method across all temporal resolutions, (b) the CompCor method at 1 s and 16 s temporal windows, and (f) the between-subject metabolic covariance estimation. In contrast, (c) the Spatiotemporal, (d) Polynomial, and (e) Baseline Normalization methods exhibit lower eigenvalue components at the highest temporal resolution (1 s), with a progressive increase as temporal resolution decreases to 16 s and 30 s. The y-axis is adjusted for each method to best display their eigenvalues.
